# An annotation dataset facilitates automatic annotation of whole-brain activity imaging of *C. elegans*

**DOI:** 10.1101/698241

**Authors:** Yu Toyoshima, Stephen Wu, Manami Kanamori, Hirofumi Sato, Moon Sun Jang, Suzu Oe, Yuko Murakami, Takayuki Teramoto, ChanHyun Park, Yuishi Iwasaki, Takeshi Ishihara, Ryo Yoshida, Yuichi Iino

## Abstract

Annotation of cell identity is an essential process in neuroscience that allows for comparing neural activities across different animals. In *C. elegans*, although unique identities have been assigned to all neurons, the number of annotatable neurons in an intact animal is limited in practice and comprehensive methods for cell annotation are required. Here we propose an efficient annotation method that can be integrated with the whole-brain imaging technique. We systematically identified neurons in the head region of 311 adult worms using 35 cell-specific promoters and created a dataset of the expression patterns and the positions of the neurons. The large positional variations illustrated the difficulty of the annotation task. We investigated multiple combinations of cell-specific promoters to tackle this problem. We also developed an automatic annotation method with human interaction functionality that facilitates annotation for whole-brain imaging.

## Introduction

Identification of the cell is an essential process in the broad fields of biology including neuroscience and developmental biology. For example, identification of cells where a gene is expressed can often be the first step in analyzing functions and interactions of the gene. Also, the identity information is required for comparing cellular activities across different animals. In order to annotate cell identities in microscopic images, features of the cells such as positions and morphologies are often compared between the samples and a reference atlas.

The nematode *C. elegans* has a unique property that all cells and their lineages have been identified in this animal (Sulston and Horvitz 1977; Sulston et al. 1983). Additionally the morphology and the connections between all 302 neurons in adult hermaphrodite were also identified by electron microscopy reconstruction (White et al. 1986). Such detailed knowledge opens up unique opportunities in neuroscience at both single-cell and network levels. Recent advances in microscopy techniques also enable whole-brain activity imaging of the worm (Schrödel et al. 2013; Prevedel et al. 2014; Kato et al. 2015; Nichols et al. 2017; Kotera et al. 2016; Tokunaga et al. 2014; Toyoshima et al. 2016; Hirose et al. 2017), even for the free-moving worms (Nguyen et al. 2016, 2017; Venkatachalam et al. 2016). The neural activities were obtained at single-cell resolution and the identities of limited numbers of neurons were annotated manually in some of the studies (Schrödel et al. 2013; Prevedel et al. 2014; Kato et al. 2015; Nichols et al. 2017; Kotera et al. 2016; Venkatachalam et al. 2016; Toyoshima et al. 2016). However, there is no systematic and comprehensive method to annotate the neurons in whole-brain activity data (Nguyen et al. 2016).

Annotation of neuronal identities in *C. elegans* is often performed based on positions of the neurons, especially for larval animals in which the neurons are located at stereotyped positions (Bargmann and Horvitz 1991; Bargmann and Avery 1995). However, for adult animals, the positions of the neurons are highly variable between animals (Nguyen et al. 2016). Several additional pieces of information can be used such as superimposed cell identity markers and morphological information of the neurons. Currently, superimposing cell identity markers, such as fluorescent proteins expressed by well-characterized cell-specific promoters, is the most popular and reliable method for neural identification. For example, Serrano-Saiz et al. 2013 showed that such methods are effective when the number of the target neurons is limited (Serrano-Saiz et al. 2013). However, integrating this approach with the whole-brain activity imaging seems difficult because it requires different markers and different fluorescent channels for every neuron in principle. Morphological information is also useful when the number of target neurons is very limited but is not readily utilized for whole-brain imaging because the neurons are distributed densely in the head region of the worms and the morphological information cannot be obtained accurately.

Several efforts for developing automatic annotation methods were reported. In order to annotate the neurons based on their positions, the information of the positions and their variations will be required. Long et al (Long et al. 2008, 2009) produced 3D digital atlas for 357 out of 558 cells from several tens of L1 animals, and related works also used the atlas (Qu et al. 2011; Kainmueller et al. 2014). The atlas consists of positions and their deviations of the cell nuclei of body wall muscles, intestine, pharyngeal neurons, and neurons posterior to the retrovesicular ganglion, as well as some other cell types. However the neurons anterior to the retrovesicular ganglion are omitted because of their dense distribution (Long et al. 2009), and the atlas cannot be applicable to the neurons in head region important for neural information processing. Aerni et al (Aerni et al. 2013) reported positions of 154 out of 959 cells from 25 adult hermaphrodites, including intestinal, muscle, and hypodermal cells, and introduced a method that integrates useful features including fluorescent landmarks and morphological information with the cell positions. Nevertheless, the positions of neurons were not reported. As far as we know, the information of the positions of the neurons in adult worms can be obtained only from the atlas produced by the EM reconstruction work (White et al. 1986). Unfortunately, the White atlas does not have the information about the variety of the positions between individual animals. Additionally, the atlas may be deformed because of inherent characteristics of the sample preparation methods for electron microscopy. Thus, experimental data of positions of neurons in adult animals were very limited. Also automatic annotation methods for neurons in the head regions of the worm have not been reported.

Here we measure the positions of the neurons in adult animals by using multiple cell-specific promoters and create a dataset. We evaluate the variations of the positions and obtain an optimal combination of the cell-specific promoters for annotation tasks based on accumulated information of cell positions. We also develop and validate an efficient annotation tool that includes both automated annotation and human interaction functionalities.

## Results

### An annotation dataset of head neurons

In this study, we focused on the head neurons of an adult animal of the soil nematode *C. elegans*, which constitute the major neuronal ensemble of this animal (White et al. 1986). The expression patterns of cell-specific promoters were used as landmarks for cell identification (Figure 1A). The fluorescent calcium indicator Yellow-Cameleon 2.60 was expressed in a cell-specific manner by using one of the cell-specific promoters and used as a fluorescent landmark. All the neuronal nuclei in these strains were visualized by the red fluorescent protein mCherry. Additionally, the animals were stained by a fluorescent dye, DiR, to label specified 12 sensory neurons following a standard method (Shaham 2006). The worms were anesthetized by sodium azide and mounted on the agar pad. The volumetric images of head region of the worm were obtained with a laser scanning confocal microscope. All the nuclei in the images were detected by our image analysis pipeline roiedit3D (Toyoshima et al. 2016) and corrected manually. The nuclei were annotated based on the expression patterns of fluorescent landmarks.

**Figure 1:**
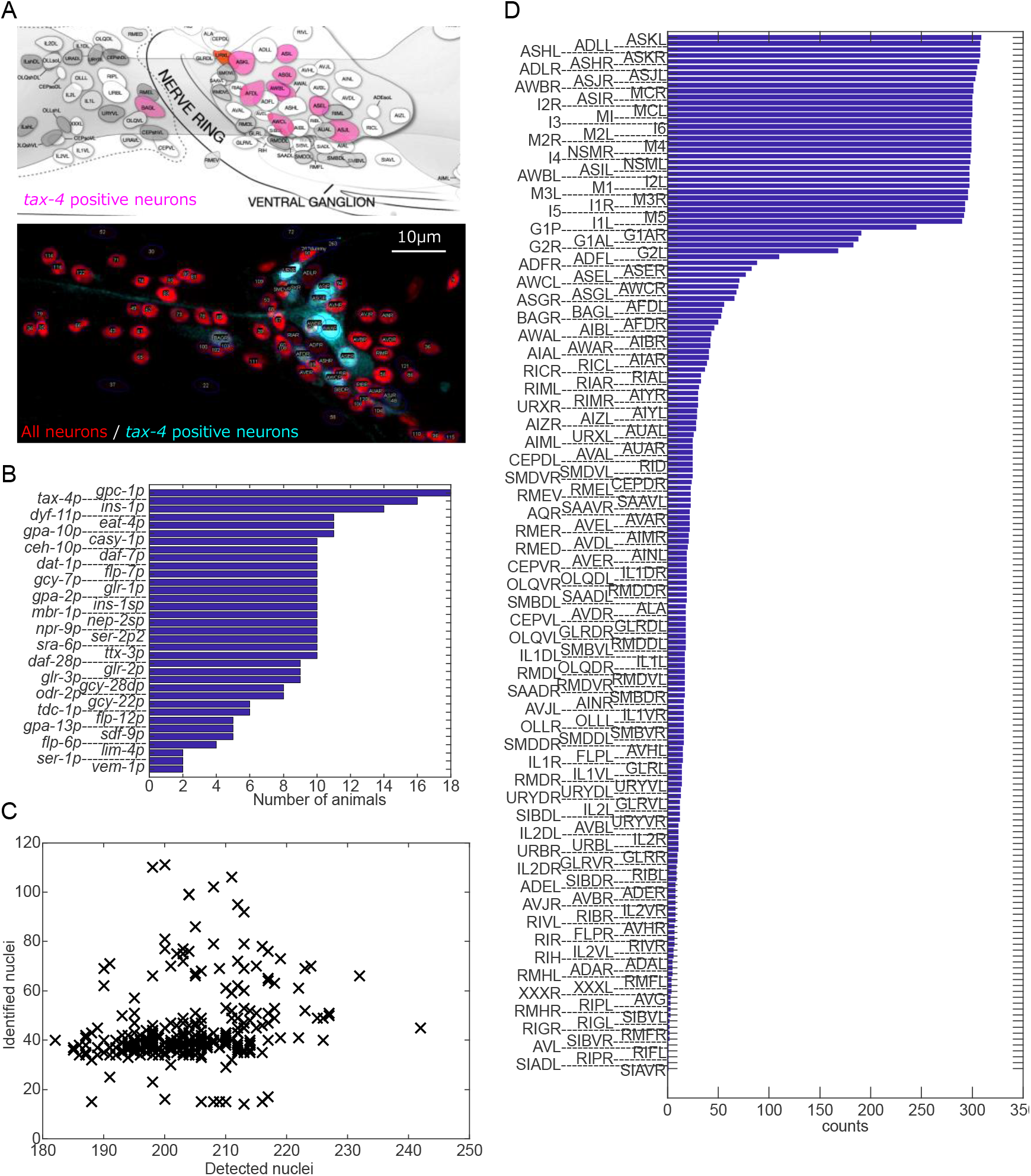
Outline of the annotation dataset. (A) The expression pattern of cell-specific promoter *tax-4p* (modified from WormAtlas) and an example image of the strain JN3006 in which the landmark fluorescent protein was expressed by *tax-4p*. Maximum intensity projection of the right side of a representative animal is shown. (B) The list of the cell-specific promoters and the number of animals used in the annotation dataset. (C) The number of the detected and the identified nuclei in each animal. (D) The names and counts of identified cells.

Finally, we obtained volumetric images of 311 animals with 35 cell-specific promoters in total (Figure 1B). On average, 203.7 nuclei were found and 44.2 nuclei were identified (Figure 1C). These positions and promoter expression information are hereafter called the annotation dataset.

Figure 1D shows names and counts of the identified cells. In most animals, 12 dye-stained cells and 25 pharyngeal cells were identified. Finally, we identified a total of 175 cells in the head region anterior to the retrovesicular ganglion. We didn’t identify URA class (4 cells), RIS cell, SIA class (2 of 4 cells) AVK class (2 cells), and RMG class (2 cells) because of the lack of suitable cell-specific promoter.

Please note that we use H20 promoter as a pan-neuronal promoter (Shioi et al. 2001). We confirmed H20 promoter was expressed in the GLR cells and XXX cells by expressing the cell-specific promoters *nep-2sp* and *sdf-9p*, respectively. We estimated that H20 promoter was expressed in pharyngeal gland cells and HMC cell, based on their positions. Also we estimated that H20 promoter was expressed weakly in the hypodermal cells, based on their positions and shape of the nuclei, but we remove these cells from our annotation dataset. We confirmed the promoter was not expressed in the socket cells nor the sheath cells by expressing the cell-specific promoter *ptr-10p*.

### Large variation disrupts position based cell annotation

How large is the variation of the relative position of the cells between individual animals? To answer this question, we need to first understand the potential sources of the variation. Intuitively, there are several possibilities: (1) placement (translational and rotational) of the worms in the obtained images, (2) curved posture of the worms (body bending), (3) inherent variation of the cell position. In order to focus on the inherent variation that we are interested in, we considered a few ways to remove the contribution of (1) and (2). PCA (principal component analysis) and subsequent alignment processes corrected the translation and rotation (see Methods). The quadratic curve fitting was employed to correct curved posture (Figure 1 - Figure Supplement 1).

After removing contribution of (1) and (2), we compiled the positions of the named cells in the annotation dataset. The positions of the nuclei identified as the same cell were collected from the annotation dataset. The mean and covariance of the positions specify a tri-variate Gaussian distribution. Three-dimensional ellipsoidal region of 2-standard deviation of the tri-variate Gaussian distribution is shown for each cell, in which about 70% of data points are expected to be included (Figure 2A). The ellipsoids largely overlap with each other, especially in the lateral ganglia (mid region of the head), because of high variation and high density of the cells. The median distance between distribution centers of neighboring cells was 4.27 μm (Figure 2B). The median length of shortest axis of the ellipsoids, equivalent to the twice of the smallest standard deviation, was about 3.78 μm. These two values were almost the same, indicating that the variation of the position of a cell reaches the mean position of the neighboring cells. Thus, the variations of the cell positions between individual animals are large.

**Figure 2:**
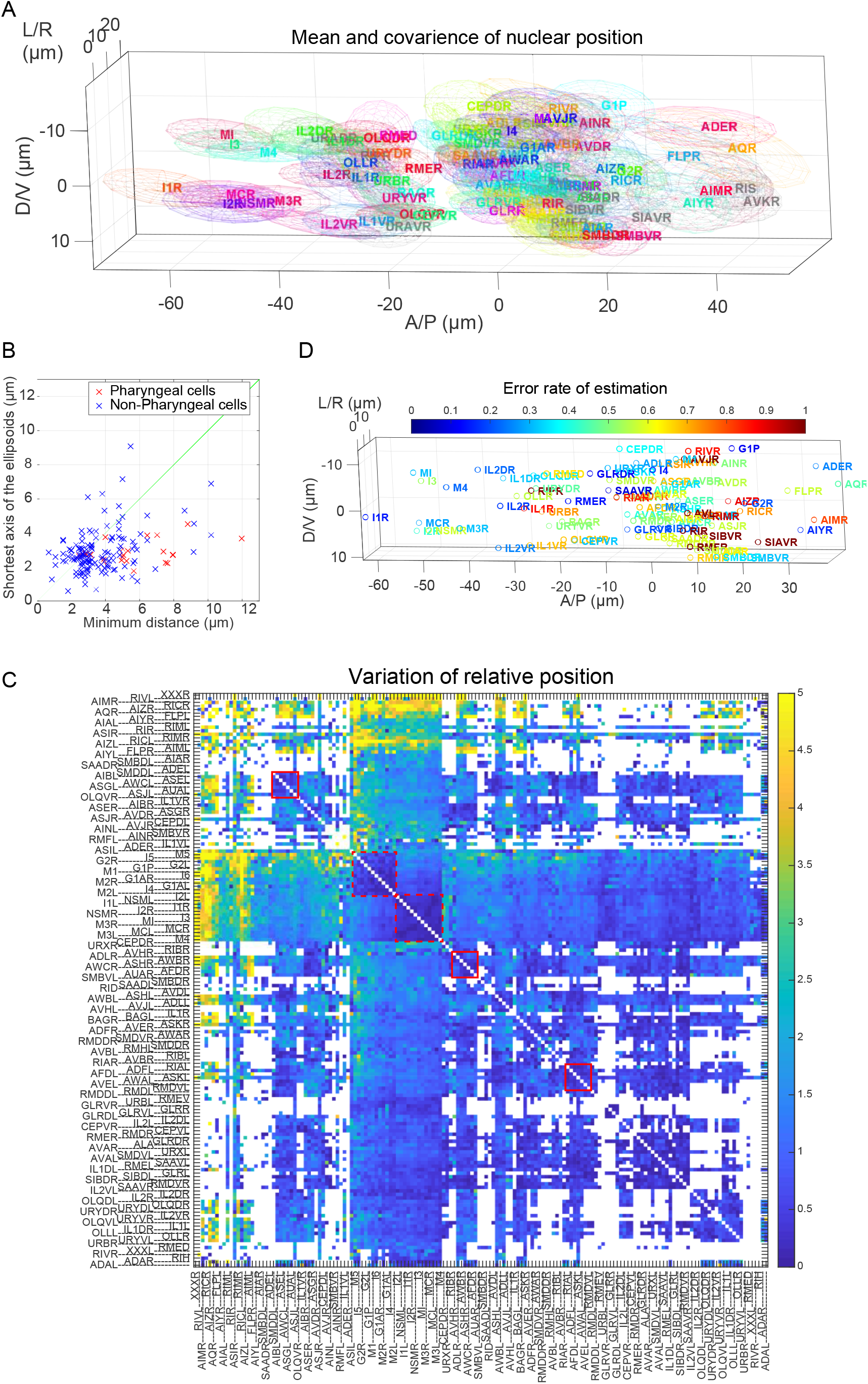
Variations of cell positions. (A) Visualization of the variation of cell positions. The ellipsoid indicates mean and covariance of the positions of the cells. Cells in the right half are shown. The colors are assigned randomly for visualization. In the case of the cells whose covariance cannot be calculated, the median of other covariance are used for visualization and shown in gray color. A/P means Anterior-posterior, D/V means Dorsal-Ventral, and L/R means Left-Right directions. (B) Minimum distance (Euclid distance of centers of nearest ellipsoids) and shortest axis length of the ellipsoids (equals to the twice of the smallest standard deviation) for each cell. The line shows where the minimum distance equals to the shortest axis length. (C) Variation of relative position of cell pairs is shown as a heat map. The red box and red dotted box indicate clusters of less varying cell pairs in lateral ganglion and pharynx, respectively. For visualization, the variations are divided by their median value, and color axis was truncated at 5 (the colors for cell pairs whose variation is larger than 5 are same as the color for cell pairs whose variation is 5). (D) The error rate of the naive estimation method is visualized with cell positions in 3D. The hot color indicates that the error rate is high.

The variations of the cell positions were explored further in a different way. We focused on the variations of the relative positions of neuron pairs. If we fix the position of a cell and align all other cells, the specific-cell-centered landscape can be drawn (Figure 2 - Figure Supplement 1). When ASKR cell was centered, the variations of positions of adjacent cells including SMDVR and ADLR were decreased, but that of other cells did not change or rather increased. When MI cell, an anterior pharyngeal neuron, was centered, the variations of positions of pharyngeal cells decreased, but those of other cells generally increased. These results suggest that the variations of relative positions are different depending on neuron pairs. We further obtained the variations of relative positions of all available cell pairs (Figure 2C). The volume of the ellipsoid of relative positions are regarded as the variation of relative positions. We found clusters of less varying cell pairs (Figure 2C, red boxes). The clusters include lateral ganglion cell pairs (Figure 2 - Figure Supplement 2) and pharyngeal cell pairs. On the other hand, there are highly varying cells including RIC, AIZ, and FLP classes.

Where do these variations of cell positions come from? In order to tackle this problem, we performed an additional analysis. Pharynx of worms often move back and forth in the body independent of other tissues. We found that the positions of dye-positive cells were affected by the positions of pharynx (Figure 2 - Figure Supplement 3); When the pharynx moved anteriorly, all the dye-positive cells (ASK, ADL, ASI, AWB, ASH, and ASJ classes) moved outside in lateral positions. In addition, anterior cells (ASK, ADL, and AWB classes) moved anteriorly, and posterior cells (ASJ class) moved posteriorly. In other words, pharynx of worms push these neurons aside. This result indicates that, although some part of the inherent variations of cell positions may come from the individual differences between animals, at least a part of the inherent variations comes from the variations of states of tissues in an animal.

How does the variations of cell positions disrupt the position-based cell annotation? Based on the mixture of the Gaussian distributions (Figure 2A), the posterior probability of assignments was calculated for the respective cells in the respective animals. The name of the cell was estimated as the name of the Gaussian which have largest probability for the cell. The error rates of this estimation method were visualized with cell positions (Figure 2D). The error rates for the cells in the posterior region were relatively low, and that for the cells in the ventral ganglion were relatively high. Mean error rate was about 50 % (see Figure 5C, described below), indicating that the variations of the cell positions actually disrupt the position-based cell annotation severely.

### Optimal combination of the cell specific promoters

In order to reduce the error rate of the annotation method, one may want to use the information of fluorescent landmarks (Kotera et al. 2016; Nguyen et al. 2016). Using multiple landmarks will reduce the error rate. One or two fluorescent channels are often available for the landmarks in addition to the channels required for the whole-brain activity imaging. We therefore sought for the optimal combination of cell-specific promoters for two-channel landmark observation using the annotation dataset.

Several properties of the promoters were evaluated in order to choose the optimal combination; how many number of cells are labelled (Figure 3A), stability of expression (Figure 3A), sparseness of the expression pattern (Figure 3B, see Methods for definition), and overlap of expression patterns in the case of combinations (See Supplementary Dataset 1). Among the 35 tested promoters, *eat-4p* was selected because it was expressed in the most numerous cells in the head region (Figure 3A). The promoters *dyf-11p* and *glr-1p* were expressed in numerous cells, and *glr-1p* was selected as the second promoter because the sparseness of the expression patterns of *glr-1p* is higher than that of *dyf-11p* (Figure 3B) and because the expression patterns of *dyf-11p* highly overlapped with that of *eat-4p*. Additionally, *ser-2p2* was selected based on the stability of the expression and low overlaps with *eat-4p* and *glr-1p*. Thus the combination of *eat-4p*, *glr-1p* and *ser-2p2* was selected (Figure 3C). The latter two promoters were used with the same fluorescent protein assuming only two fluorescent channels can be used for the landmarks as is the case for our experimental setup for whole-brain imaging. In the annotation dataset, *eat-4p* was expressed in 69 cells and *glr-1p* + *ser-2p2* were expressed in 50 cells out of 196 cells in the head region of adult worms.

**Figure 3:**
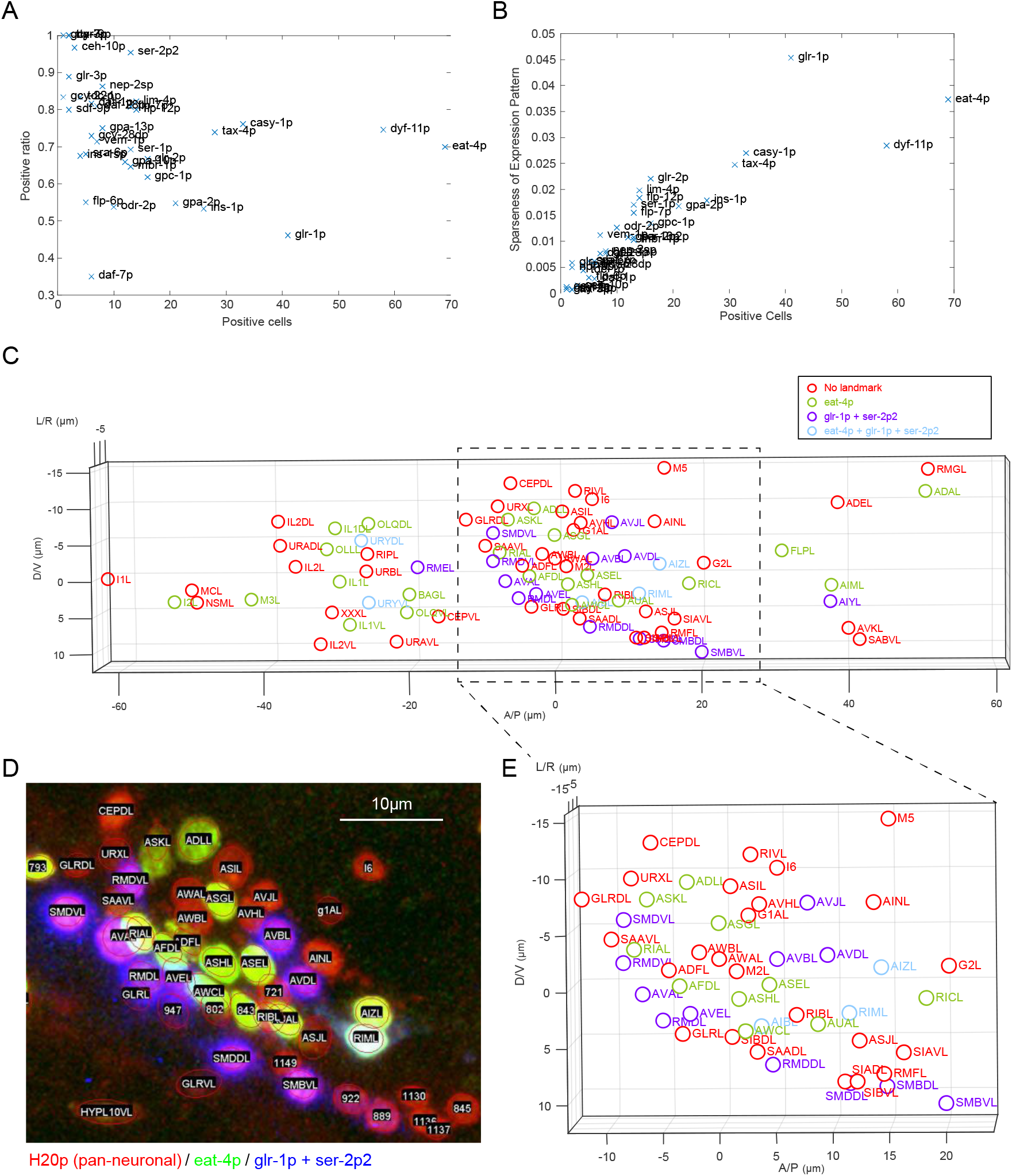
Optimal combination of the cell-specific promoters. (A) Number of positive cells and mean positive ratio of the cell-specific promoters. (B) Number of positive cells and sparseness of expression pattern of the cell-specific promoters. (C) Visualization of the optimal combination of the cell-specific promoters. The cells in the right half are shown. (D) An example fluorescent image of JN3039 strain. (E) A part of (C) was zoomed for comparison with (D).

All combinations of the promoters could be evaluated by an algorithm that considers the number of expression, sparseness and overlap of expression patterns (see Methods and Supplementary Table 1). In brief, the algorithm highly evaluated a combination when two neighboring cells were in different colors. In the case of three promoters and two fluorescent channels, the combination consisting of *eat-4p*, *glr-1p* and *ser-2p2* was placed in the 18th rank out of the possible 20825 combinations.

Here we produced a strain JN3039 as follows. The far-red fluorescent protein tagRFP675 was expressed using *eat-4p*, and the blue fluorescent protein tagBFP was expressed using *glr-1p* and *ser-2p2*. The red fluorescent protein mCherry was expressed using the pan-neuronal promoter H20p. This strain does not use fluorescent channels of CFP, GFP, and YFP and is useful for the cell identification tasks. For example, if there is a strain that express these fluorescent proteins with a promoter whose expression patterns should be identified, one can do it by just crossing the strain with our standard strain JN3039.

Additionally a strain JN3038 was made from the strain JN3039 by expressing fluorescent calcium indicator Yellow-Cameleon 2.60 with the pan-neuronal promoter H20p. This five-colored strain will enable whole-brain activity imaging with annotation.

### Generating atlases for automatic annotation

We confirmed that the positions of the cells in the worm show so large variations that using a single reference atlas for annotation is not sufficient. Instead, we propose to generate a large set of atlases that captures such positional variations using the 311 partially annotated data in order to exploit statistical methods, such as majority voting (to be illustrated in the next section), for a more effective automatic annotation.

To obtain an atlas with fully annotated cells, we need to combine positional information of cells from multiple partially annotated animals while maintaining the relative position between the cells as much as possible. We assembled the partially identified cell positions in our annotation dataset as follows (Figure 4A, see Methods for detail):

A. Choose an animal 1 and extract the positions of the identified cells.
B. Choose an animal 2 and register it to animal 1 based on the positions of commonly-identified cells in both animals.
C. Map the cells in animal 2 that were not identified in the animal 1 according to the registration, and add them to the identified cell list of animal 1.
D. Repeat steps B) and C) until all the animals were covered.
E. Add the positions of not-identified cells in our annotation dataset using the positions of the cells in White’s atlas.

The resulting atlas depends on the order of assembly, which reflects the variation of the cell positions between individual animals. By random sampling of the animals, we generated around 3000 synthetic atlases that are used to reduce the error rate of the estimation.

We compared the position of cells in the atlases and the dataset. We visualized mean and covariance of the positions of the cells in the atlases (Figure 4B). The outline of the cell positions in the atlas was similar to that in the dataset, suggesting that our atlases capture the positional variations of the cells in the dataset. The distances of mean positions of the cells between the atlases and dataset was small (Figure 4C, 1.43 μm in median), but become large for the rarely detected cells. Volumes of the ellipsoids (covariance of the positions of the cell) between the atlas and the dataset were also similar (Figure 4D), but the volume for the atlases was larger than that for the dataset when the cell was rarely detected in the dataset. The variation of relative positions of the cell pairs in the atlases was similar to that in the dataset when the cell-pair was co-detected in the dataset frequently enough, and become larger when the cell pair was less co-detected (Figure 4E and Figure 4 - Figure Supplement 1). These results indicate that our atlases capture the positional variations of the cells, and the atlases will be more precise when the dataset includes a lot more animals and annotated cells.

### Automatic annotation using bipartite matching and majority voting

Having a set of atlases that capture the positional variations of the cells, we can account for the spatial uncertainty of cell annotation using majority voting. We propose an automatic annotation method that utilizes bipartite matching and majority voting (Figure 5A). Each part will be described below in brief (see Method for detail).

In the bipartite matching step, the cells in a target animal were assigned to the cells in an atlas. An assignment of a cell in the target animal to a cell in the atlas has a cost based on the similarity (or dissimilarity) of the two cells including Euclidean distance, expressions of landmark promoters, and the feedback from human annotation. The optimal combination of the assignments that minimizes the sum of the costs was obtained by using Hungarian algorithm. The name of the cell in the target animal can be estimated as the name of the assigned cell in the atlas in this step.

To handle the configurational variations of cells to be annotated, we used the majority voting technique. Assuming the generated atlases could capture the positional variations of the cells, we assigned unannotated cells in the target animal to those in *N*_*a*_ atlases, then giving *N*_*a*_ annotation results. Each assignment of a cell is considered as a vote, and the most voted assignment was considered as the top rank estimation of annotation.

In order to validate our automatic annotation method, a 5-fold cross validation test was performed. All the animals in the annotation data set were randomly divided into five subsets. We perform a total of 5 tests. For each test, we exclude one of the subset from training of atlases, and use it to estimate the annotation performance based on the trained atlases. The error rate of bipartite matching was relatively high, and the majority voting could deliver significant improvements of the annotation accuracy (Figure 5B). On average, 78.3 nuclei were annotated and 46.2 nuclei were successfully estimated as the top rank, and the error rates of the top rank estimation was 41.4% (Figure 5B and C). As a control, two methods are introduced; one method only considers the mean and covariance of the cell positions of raw data (without using the atlases and voting, see Figure 2D). The other method considers the mean and covariance of the cell positions in the atlases (without using majority voting). The error rate of the two methods were higher than the proposed method, indicating that the majority voting step in the proposed method contribute to the correct estimation. If we consider the accuracy for the top 5 voted estimations (shown as rank 5), the error rate decreased to 7.3%.

The automatic annotation method was applied to the animals with fluorescent landmarks (strain JN3039, see Figure 3C-E). With the help of the optimized expression of landmark fluorescent proteins, the number of identified cells in an animal will increase compared to the strains used to make the annotation dataset. On average, 202 nuclei were found and 156.3 nuclei were identified from 15 adult animals. The error rates of the top rank estimation with and without fluorescent landmark were 37.7% and 51.5%, respectively, indicating that utilizing the fluorescent landmark also contribute the correct estimation (Figure 5D). If we consider the accuracy for the top 5 voted estimations, the error rate decreased to 8.1%. These error rates were comparable to the cross-validation results for the annotation dataset, suggesting that our annotation framework will work correctly for the whole-brain activity imaging.

The automatic annotation method was also applied to the animals in a microfluidic chip for whole-brain activity imaging (Figure 5 - figure supplement 1). The error rates of the top rank estimation with and without fluorescent landmark were 52.1% and 72.8%, respectively, and that of the top 5 voted estimations was 12.2%. The worms were compressed and distorted to be held in the microfluidic chips, and the distortion of the worm may increase the error rates. During whole-brain imaging for free-moving animals (Nguyen et al. 2016, 2017; Venkatachalam et al. 2016), the worms will be less compressed and less distorted, and our algorithm may works better.

Additionally, our algorithm is implemented in the GUI roiedit3d (Toyoshima et al. 2016), and it can handle feedback information from the human annotations. Once annotations are corrected manually, our method can accept corrections and uses them to improve the results. For example, one can identify neurons manually by using other information including the neural activity or morphology, and the automatic estimation for the other neurons will be improved. The final results can be added to the annotation dataset and the annotation algorithm will work more accurately. Thus the feedback system incorporates tacit knowledge into the automatic annotation method. Through the interactive process our algorithm will make human annotation tasks more efficient.

## Discussion

In this study, we obtained volumetric fluorescent image of 311 animals using 35 promoters, and created an annotation dataset that contains the positions of the identified cells and expression patterns of promoters in respective animals. Utilizing the annotation dataset we evaluate the variation of the positions of the cells and choose the combination of the promoters optimal for our annotation tasks. We proposed an automatic annotation method and validated its performance on head neurons of adult worms for whole-brain imaging. Thus, we successfully integrate the annotation techniques with the whole-brain activity imaging.

The cell positions of real animals and its variation will be the most important information for the cell identification. As far as we know, this might be the first report about the large-scale information of the positions of the cells in the head region of adult *C. elegans*, which lead to systematic and comprehensive method to annotation of the head neurons. The error rate of the automatic annotation might be slightly high for fully automatic annotation. The integration of the automatic annotation method to the GUI enables machine-assisted annotation and enhances the process of whole-brain image annotation. Increasing the number of animals and promoters will improve the accuracy and objectivity of the automatic annotation method.

Increasing the number of fluorescent channels and landmarks will also improve the accuracy. Long-stokes shift fluorescent proteins might be good candidates because they use irregular fluorescent channels that will not be used in standard application. In our case, however, these proteins disrupted the neighbor fluorescent channels by leaking-out. Employing color deconvolution techniques will increase the number of substantial fluorescence channels and may improve the accuracy.

The images of the animals we recorded will have useful information for annotation including size of the nuclei and intensities of the fluorescence. In the manual annotation process we utilized these pieces of information for improving accuracy. On the other hand our automatic annotation algorithm does not utilizes these pieces of information and it may be one of the causes of relatively low accuracy of the algorithm. Recent advances in artificial neural networks especially in the field of image analysis will enable to utilize such information for automatic annotation. It is well known that artificial neural networks require large amount of training data composed of images and the corresponding grand truth. Our annotation dataset contains images with identity information and will be ideal for the training data, but the number of data may not be enough. Our method that makes annotation more efficient will play an important role for opening up the path to utilization of artificial neural networks in the future.

There are no dataset of cell positions that can be used as a benchmark of cell identification methods. For example, a new method that solve the cell identification problem as a nonlinear assignment problem was reported recently (Bubnis et al. 2019). The report utilizes synthesized data and does not use real data. To evaluate the real performance of new methods, the method should be tested on real data. Our annotation dataset will be an ideal benchmark of newly developed cell identification methods. Thus our study will facilitate the future studies for automatic annotation methods.

In order to identify the expression patterns of the promoters, the most accurate method is testing whether the fluorescence of promoter overlaps with the fluorescence of the neuronal identity markers (Serrano-Saiz et al. 2013). In such cases our standard strain and automatic annotation method will help the selection of the markers through objective estimation of cell identities.

Our framework of creating the annotation dataset and developing automatic annotation method can be applied to species other than *C. elegans*. For covering all neurons, the number of available cell-type specific promoters and their variety will be important.

## Methods

### Strains and cultures

*C. elegans* strains used in this study are listed in Table 1. Animals were raised on nematode growth medium at 20°C. E. coli strain OP50 was used as a food source.

**Table 1:**
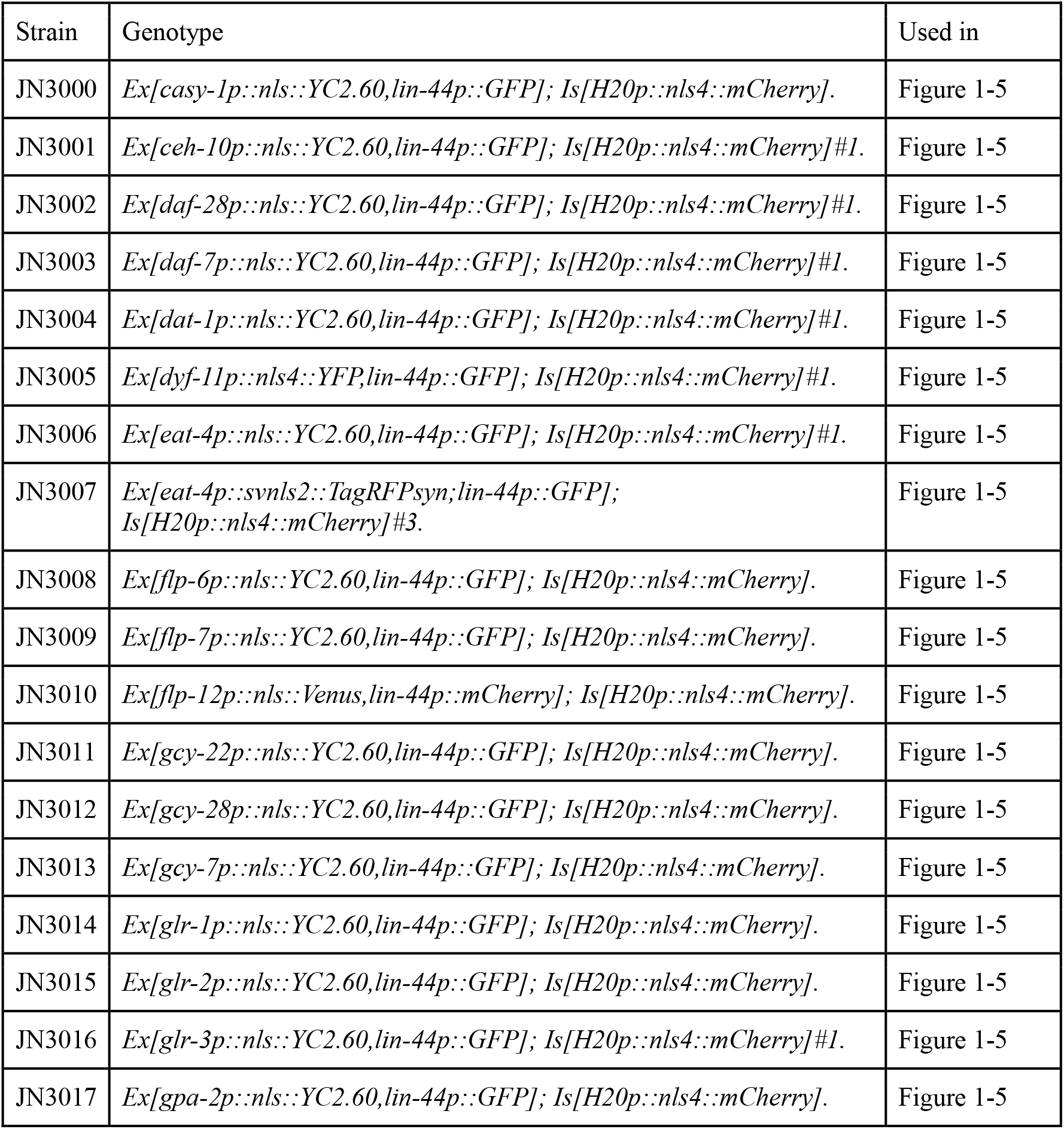
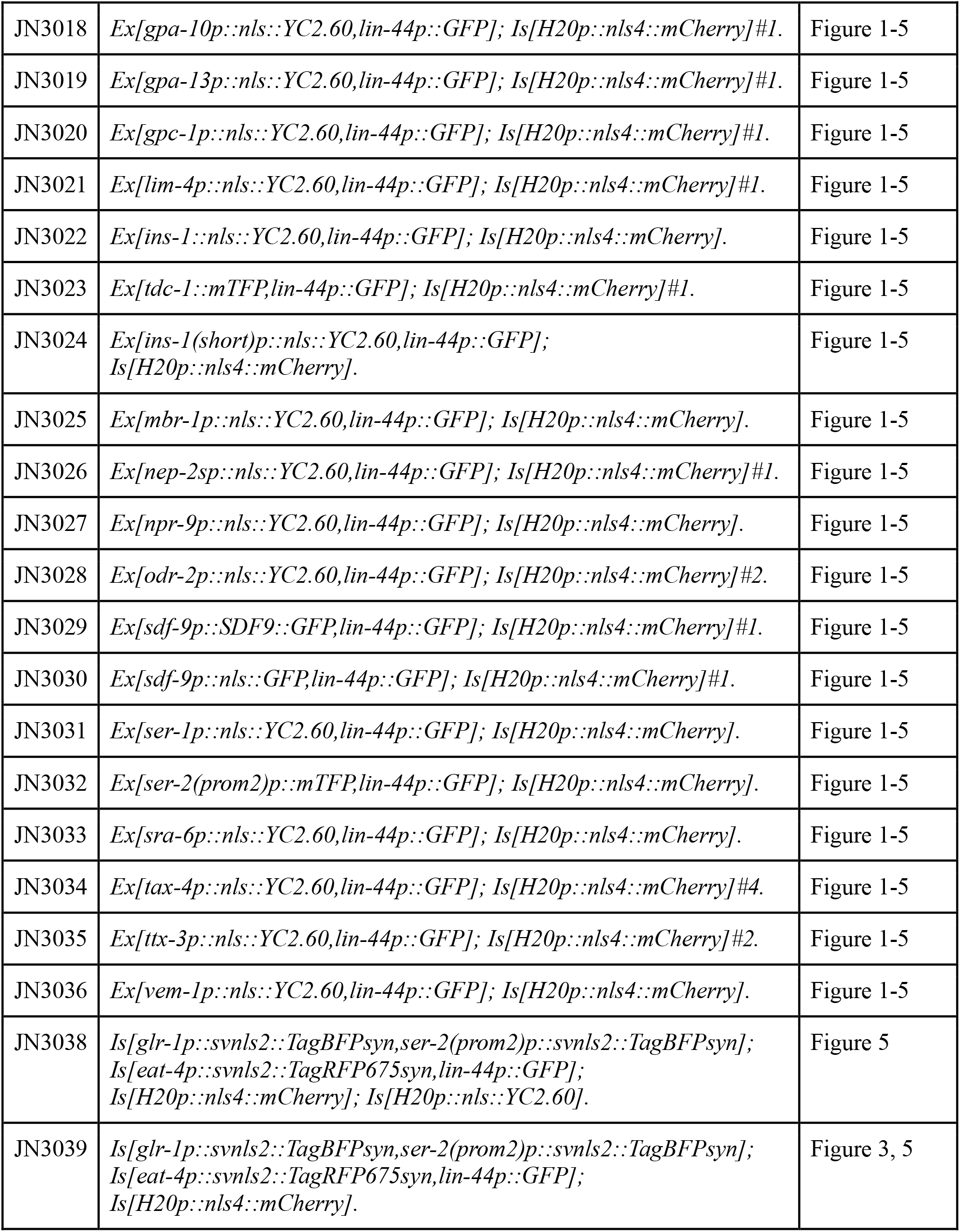
Strain list used in this study.

| Strain | Genotype | Used in |
| --- | --- | --- |
| JN3000 | *Ex[casy-1p::nls::YC2.60,lin-44p::GFP]; Is[H20p::nls4::mCherry].* | Figure 1-5 |
| JN3001 | *Ex[ceh-10p::nls::YC2.60,lin-44p::GFP]; Is[H20p::nls4::mCherry]#1.* | Figure 1-5 |
| JN3002 | *Ex[daf-28p::nls::YC2.60,lin-44p::GFP]; Is[H20p::nls4::mCherry]#1.* | Figure 1-5 |
| JN3003 | *Ex[daf-7p::nls::YC2.60,lin-44p::GFP]; Is[H20p::nls4::mCherry]#1.* | Figure 1-5 |
| JN3004 | *Ex[dat-1p::nls::YC2.60,lin-44p::GFP]; Is[H20p::nls4::mCherry]#1.* | Figure 1-5 |
| JN3005 | *Ex[dyf-11p::nls4::YFP,lin-44p::GFP]; Is[H20p::nls4::mCherry]#1.* | Figure 1-5 |
| JN3006 | *Ex[eat-4p::nls::YC2.60,lin-44p::GFP]; Is[H20p::nls4::mCherry]#1.* | Figure 1-5 |
| JN3007 | *Ex[eat-4p::svnls2::TagRFPsyn;lin-44p::GFP]; Is[H20p::nls4::mCherry]#3.* | Figure 1-5 |
| JN3008 | *Ex[flp-6p::nls::YC2.60,lin-44p::GFP]; Is[H20p::nls4::mCherry].* | Figure 1-5 |
| JN3009 | *Ex[flp-7p::nls::YC2.60,lin-44p::GFP]; Is[H20p::nls4::mCherry].* | Figure 1-5 |
| JN3010 | *Ex[flp-12p::nls::Venus,lin-44p::mCherry]; Is[H20p::nls4::mCherry].* | Figure 1-5 |
| JN3011 | *Ex[gcy-22p::nls::YC2.60,lin-44p::GFP]; Is[H20p::nls4::mCherry].* | Figure 1-5 |
| JN3012 | *Ex[gcy-28p::nls::YC2.60,lin-44p::GFP]; Is[H20p::nls4::mCherry].* | Figure 1-5 |
| JN3013 | *Ex[gcy-7p::nls::YC2.60,lin-44p::GFP]; Is[H20p::nls4::mCherry].* | Figure 1-5 |
| JN3014 | *Ex[glr-1p::nls::YC2.60,lin-44p::GFP]; Is[H20p::nls4::mCherry].* | Figure 1-5 |
| JN3015 | *Ex[glr-2p::nls::YC2.60,lin-44p::GFP]; Is[H20p::nls4::mCherry].* | Figure 1-5 |
| JN3016 | *Ex[glr-3p::nls::YC2.60,lin-44p::GFP]; Is[H20p::nls4::mCherry]#1.* | Figure 1-5 |
| JN3017 | *Ex[gpa-2p::nls::YC2.60,lin-44p::GFP]; Is[H20p::nls4::mCherry].* | Figure 1-5 |
| JN3018 | *Ex[gpa-10p::nls::YC2.60,lin-44p::GFP]; Is[H20p::nls4::mCherry]#1.* | Figure 1-5 |
| JN3019 | *Ex[gpa-13p::nls::YC2.60,lin-44p::GFP]; Is[H20p::nls4::mCherry]#1.* | Figure 1-5 |
| JN3020 | *Ex[gpc-1p::nls::YC2.60,lin-44p::GFP]; Is[H20p::nls4::mCherry]#1.* | Figure 1-5 |
| JN3021 | *Ex[lim-4p::nls::YC2.60,lin-44p::GFP]; Is[H20p::nls4::mCherry]#1.* | Figure 1-5 |
| JN3022 | *Ex[ins-1::nls::YC2.60,lin-44p::GFP]; Is[H20p::nls4::mCherry].* | Figure 1-5 |
| JN3023 | *Ex[tdc-1::mTFP,lin-44p::GFP]; Is[H20p::nls4::mCherry]#1.* | Figure 1-5 |
| JN3024 | *Ex[ins-1(short)p::nls::YC2.60,lin-44p::GFP]; Is[H20p::nls4::mCherry].* | Figure 1-5 |
| JN3025 | *Ex[mbr-1p::nls::YC2.60,lin-44p::GFP]; Is[H20p::nls4::mCherry].* | Figure 1-5 |
| JN3026 | *Ex[nep-2sp::nls::YC2.60,lin-44p::GFP]; Is[H20p::nls4::mCherry]#1.* | Figure 1-5 |
| JN3027 | *Ex[npr-9p::nls::YC2.60,lin-44p::GFP]; Is[H20p::nls4::mCherry].* | Figure 1-5 |
| JN3028 | *Ex[odr-2p::nls::YC2.60,lin-44p::GFP]; Is[H20p::nls4::mCherry]#2.* | Figure 1-5 |
| JN3029 | *Ex[sdf-9p::SDF9::GFP,lin-44p::GFP]; Is[H20p::nls4::mCherry]#1.* | Figure 1-5 |
| JN3030 | *Ex[sdf-9p::nls::GFP,lin-44p::GFP]; Is[H20p::nls4::mCherry]#1.* | Figure 1-5 |
| JN3031 | *Ex[ser-1p::nls::YC2.60,lin-44p::GFP]; Is[H20p::nls4::mCherry].* | Figure 1-5 |
| JN3032 | *Ex[ser-2(prom2)p::mTFP,lin-44p::GFP]; Is[H20p::nls4::mCherry].* | Figure 1-5 |
| JN3033 | *Ex[sra-6p::nls::YC2.60,lin-44p::GFP]; Is[H20p::nls4::mCherry].* | Figure 1-5 |
| JN3034 | *Ex[tax-4p::nls::YC2.60,lin-44p::GFP]; Is[H20p::nls4::mCherry]#4.* | Figure 1-5 |
| JN3035 | *Ex[ttx-3p::nls::YC2.60,lin-44p::GFP]; Is[H20p::nls4::mCherry]#2.* | Figure 1-5 |
| JN3036 | *Ex[vem-1p::nls::YC2.60,lin-44p::GFP]; Is[H20p::nls4::mCherry].* | Figure 1-5 |
| JN3038 | *Is[glr-1p::svnls2::TagBFPsyn,ser-2(prom2)p::svnls2::TagBFPsyn]; Is[eat-4p::svnls2::TagRFP675syn,lin-44p::GFP]; Is[H20p::nls4::mCherry]; Is[H20p::nls::YC2.60].* | Figure 5 |
| JN3039 | *Is[glr-1p::svnls2::TagBFPsyn,ser-2(prom2)p::svnls2::TagBFPsyn]; Is[eat-4p::svnls2::TagRFP675syn,lin-44p::GFP]; Is[H20p::nls4::mCherry].* | Figure 3, 5 |

### Microscopy

A set of static 3D multi-channel images of *C. elegans* strains ranging from JN3000 to JN3036 were obtained as follows. Day 1 adult animals were stained by the fluorescent dye DiR (D12731, Thermo Fisher Scientific) with the standard method (Shaham 2006). The stained animals were mounted on a 2% agar pad and paralyzed by sodium azide. The fluorescence of the fluorescence proteins and the dye was observed sequentially using laser scanning confocal microscopy (Leica SP5 with 63× water immersion lens and 2× zoom). The sizes of the images along the x1 and x2 axes were 512 and 256 voxels, respectively, and the size along the x3 axis varied depending on the diameter of the animal. The sizes of a voxel along the x1, x2, and x3 axes were 0.240, 0.240, and 0.252 μm, respectively.

A set of 3D multi-channel images of strain JN3039 was obtained as described above without using the fluorescent dye DiR.

A set of 3D multi-channel images of strain JN3038 was obtained as follows. Day 1 adult animals were conditioned on NGM plate with OP50 (Kunitomo et al. 2013). The conditioned animals were introduced and held in a microfluidic device called olfactory chip (Chronis, Zimmer, and Bargmann 2007). The depth and width of the fluid channel in the chip were modified in order to reduce the distortion of the worms. The animals and their head neurons moved to some extent in the device because the animals were not paralyzed. The fluorescence of the tagBFP, tagRFP675, and mCherry channels was observed simultaneously using customized spinning disk confocal microscopy. The sizes of the image along the x1 and x2, and x3 axes were 512, 256, and 50 voxels, respectively. The sizes of a voxel along the x1, x2, and x3 axes were 0.28, 0.28, and about 0.77 μm, respectively.

### Image analysis for the annotation dataset

All the nuclei in the images were detected by our image analysis pipeline roiedit3D (Toyoshima et al. 2016) and corrected manually. The cells stained by the chemical dye were identified as reported (Shaham 2006). The cells marked by cell-specific promoters were identified based on the reported expression patterns and positions of the nuclei. The nuclei of the pharyngeal cells were also identified based on the positions of the nuclei.

### Correction of posture of worms

First, all the positions of nuclei in a worm determined by roiedit3D were analyzed by PCA and the 1st principal component axis (PC1 axis) were defined as the anterior-posterior axis. The positions of the nuclei were fitted with a quadratic function along the PC1 axis (see Figure 1 - Figure supplement 1). The determined quadratic function minimizes the sum of the squared distances from the fitted line to the positions of nuclei along PC2-PC3 axis. The positions were corrected so that the quadratic line was straightened and at the same time the rolled posture of the animal was corrected. The positions of the nuclei were projected onto the plane with PC2-PC3 axes and the sparsest direction from the center was defined as dorsal direction. The positions were rotated along the PC1 axis so that PC1 (antero-posterior axis) is aligned to x axis the dorsal direction is aligned to positive direction of y axisq. Then we estimated the anterior direction based on the density of the lateral cells. The densest position was set as the origin of the anterior-posterior axis. The origins of the dorsal-ventral and left-right axes were the same as the origin of the PC2 and PC3 axes. The worms can be aligned by these procedures. The positions of the animals in the annotation dataset were corrected precisely based on the positions of the dye-stained cells.

### Variation of relative positions

Variation of relative position of a cell pair was calculated as the determinant of the covariance of relative cell positions. Let *X*_*i*_ and *Y*_*i*_ be the position of the cell X and Y in the i-th animal, respectively, and the cells were identified in *n* animals.

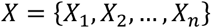

and

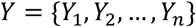

are n-by-3 matrices of the positions of the cells X and Y, respectively. Then the variation of relative positions of cell X and cell Y is

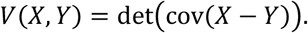

For visualization, *V*(*X*, *Y*) was divided by median value of all *V*. The pairs with *n* ≤ 3 were ignored because the determinant of covariance cannot be calculated.

Less varying cell pairs were found based on permutation of animals (permutation test). A permutation of the vector *X* permutes the order of elements of the vector *X*, for example,

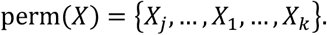

The pair of cell X and Y was regarded as less varying if

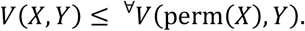

For the pairs of 4 ≤ *n* ≤ 10, all the permutations (equal to or less than 10! ~ 3.6 × 10^6^ combinations) were calculated. For the pairs of *n* > 10, 1 × 10^7^ permutations were randomly selected and calculated.

### The algorithm for searching optimal combination of cell-specific promoters and the definition of the sparseness

The most important factor for selecting promoters in order to improve annotation accuracy is to achieve a checkerboard-like coloring pattern for the ease of separating neighboring cells. A simple metric to account for this factor is to sum the number of neighboring cell pairs that exhibit a different color based on cell-specific promoters, where each pair is inversely weighted by the distance between the two neurons. Such a metric can be considered as a modification to an Ising model in physics. We choose a Gaussian probability model for the weighting function with an empirically chosen value of the standard deviation to be 9.6 μm. The metric *M* can be written as

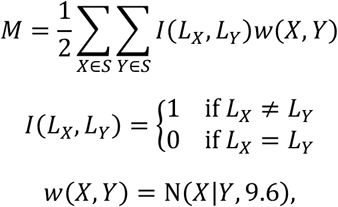

where *S* is a set of all cells in an animal. *X* and *Y* are positions of cell X and cell Y, respectively. *L*_*x*_ is label of cell X and *L*_*x*_ = (1,0) means that landmark protein of color 1 is expressed in the cell X but that of color 2 is not expressed. Because the experimental setup has a limited amount of channels, we are able to perform an exhaustive search for all possible combination of the available promoters, and compare the final values of the metric as a reference for choosing the combination of cell-specific promoters used in our experiment. We evaluated all the combinations for 3 promoters and 2 colors (20825 combinations). The scores of the single promoter for single color were used as the index of sparseness.

### Generating atlases

To obtain an atlas with fully annotated cells, we need to combine positional information of cells from multiple partially annotated images while maintaining the relative position between the cells as much as possible. We achieve this goal by maximizing the consistency (or smoothness) of a displacement flow when combining different images, for which the displacement flow is defined as follows.

Suppose that in two images, denoted by *I*_0_ and *I*_1_, there coexist *C* annotated cells. The displacement of cell *i* is denoted by  where 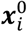 and 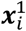 denote the positions of the cell in *I*_0_ and *I*_1_, respectively. Then, we define a displacement flow field ***d***_0→1_(***x***) from *I*_0_ to *I*_1_ on the entire space ***x*** ∈ ℝ^3^:

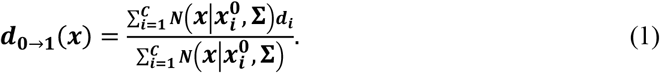

Here, *N*(***x***|***μ***, *Σ*) denotes the density function of the normal distribution with mean ***μ*** and covariance Σ (Please note that *Σ* is 3 × 3 covariance matrix and determine effective range of the displacement of a cell). This represents a flow field function interpolated by the given displacements of the *C* cells in the two images. When taking the weighted average in the calculation of ***d***_0→1_(***x***), larger weights are assigned to the displacements of more neighboring cells with respect to ***x*** in *I*_0_.

To generate an atlas, we conducted the following steps (see Figure 4A):

1. Set a randomly ordered sequence {*I*_1_,…, *I*_311_} of the 311 partially annotated animals. We discard the sequence if the *I*_1_ has less than 60 annotated cells.
2. For *t* ∈ {2 …, 311}, cells in *I*_*t*_ were sequentially aligned to those in *I*_1_ as follows:

A. The positions of all annotated cells in *I*_1_ were unchanged.
B. All annotated cells that coexisted in both *I*_1_ and *I*_*t*_ were used to calculate the displacement field ***d***_*t*→1_(***x***) with a pre-determined *Σ* (Eq 1).
C. All cells annotated in *I*_*t*_ but not in *I*_1_ with their positions denoted by ***x***_*t*_ were shifted and aligned to *I*_1_ according to ***x***_1_ ← ***x***_*t*_ + ***d***_*t*→1_(***x***_*t*_). Add them to the annotated cells in *I*_1_.
D. Terminate the iteration if all annotated cells have been aligned in the synthesized reference image.

**Figure 4:**
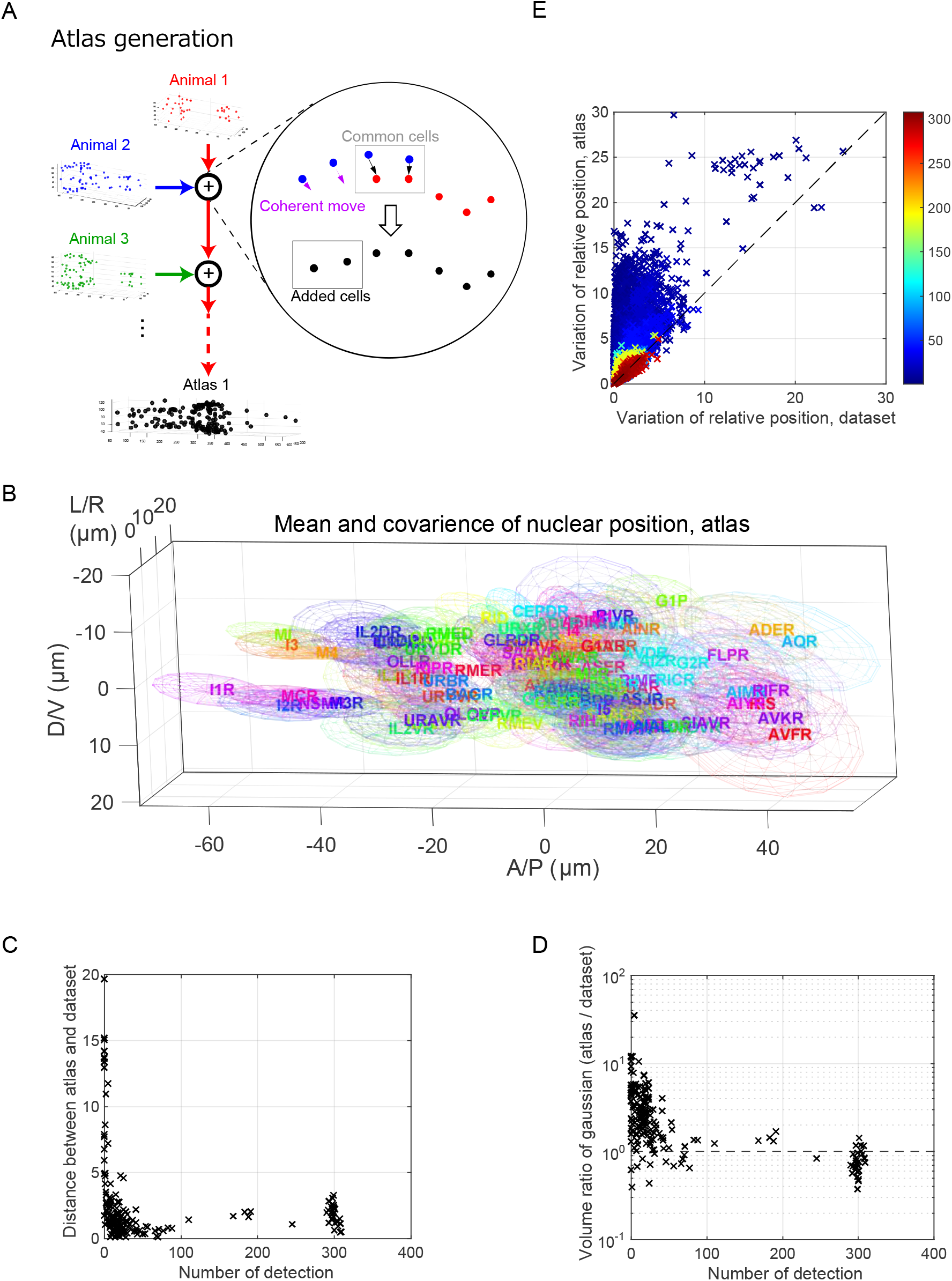
Atlas generation. (A) The outline of the atlas generation method. (B) Visualization of the variation of the cell positions in synthetic atlas. The cells and colors are same as Figure 2A. (C) Distance of mean position of cells between the atlas and the dataset. (D) Ratio of volume of ellipsoid (covariance of the positions of the cell) between the atlas and the dataset. (E) Comparing the variation of relative positions of dataset and that of atlas. The color indicates how many times the neuron pair is co-detected in an animal of the dataset.

In this scheme, a spatial pattern of produced cells was largely affected by the interpolated flow fields. In general, the performance will be poor if the number of observed source displacements was small. To reduce such instability, we skipped *I*_*t*_ and used it later when the *I*_*t*_ shared less than half cells annotated in common with *I*_1_. Repeating this procedure, we generated 3,000 reference samples.

The generated reference samples serve as a set of virtual atlases that imitate observed topological variations of cellular positions across different worm samples. To obtain more realistic atlases, we optimized *Σ* = diag(*σ*_1_, *σ*_2_, *σ*_3_) in Eq 1, which is the parameter to control the smoothness of displacements in the sequential alignments. We defined an objective function to reflect the similarity of the topological variations between our raw data set and the generated atlas. By optimizing such objective function and taking the optimal values of the parameters as a reference, we selected an empirical value of *Σ* = diag(9.6 μm, 9.6 μm, 9.6 μm). Details of the objective function and optimization is in Supplementary Note 1.

### Bipartite graph matching

Detected cells in a target animal and an atlas were matched using the Hungarian algorithm to solve the bipartite graph matching problem. The matching was achieved by comparing one or more selected features between cells. Here, features refer to some quantitative properties for the cells that can be used to distinguish the identity of a cell from another. The most fundamental feature is the positions of cells. Other typical features include cell volume, fluorescence intensities, and so on. We use expression of landmark proteins (i.e. binarized fluorescent intensities) and feedback from human annotation. With such features, the dissimilarity of cells was represented by a matrix *A*, where the {*i*, *j*} entry is the distance of the feature values between the *i*^th^ cell in the target and the *j*^th^ cell in the atlas. When there are *N*_*f*_ features chosen, we can assemble them into a single matrix *A*_BGM_ through a weighted sum:

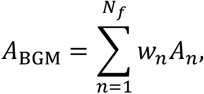

where *w*_*n*_ is the weight for each feature. We use *w*_*distance*_ = 1, *w*_*landmark*_ = 20. For feedback from human annotation, the assignments incompatible with the human annotation have infinity dissimilarity. With a given assignment, we can calculate the sum of the dissimilarity values in *A*_BGM_ that correspond to the selected matching. A modified Hungarian algorithm (Jonker and Volgenant 1987) was used to minimize the total distance with respect to all possible assignments under the constraint of one-to-one matching.

### Majority voting

Multiple name assignments of a cell in the subjective animal were obtained by repeating the bipartite graph matching using 500 different atlases. Each assignment was considered as one vote, and the estimated names for a target cell was ranked by vote counts. The estimation for a cell was independent of each other and multiple cells may have the same estimated names. If non-overlapping result is required, one can assemble cost matrix based on vote counts and apply the Hungarian algorithm.

### Calculation of error rate of automatic annotation

All the detected cells in a target animal other than hypodermal cells were used as a target. The names of the cells were estimated by our automatic annotation method based on their positions. Expression of landmark promoters were also used for Figure 5D and figure 5 Supplementary figure 1. The estimated results are compared to the human annotation (grand truth). Our automatic annotation method returns multiple ranked candidates for a target cell. The rank N error rate indicates that it is considered correct if the correct annotation appeared in the top N estimations. Un-annotated cells were ignored in calculating error rate. The animals that have less annotated cells were removed to avoid the effect of deviation of the annotated cells.

**Figure 5:**
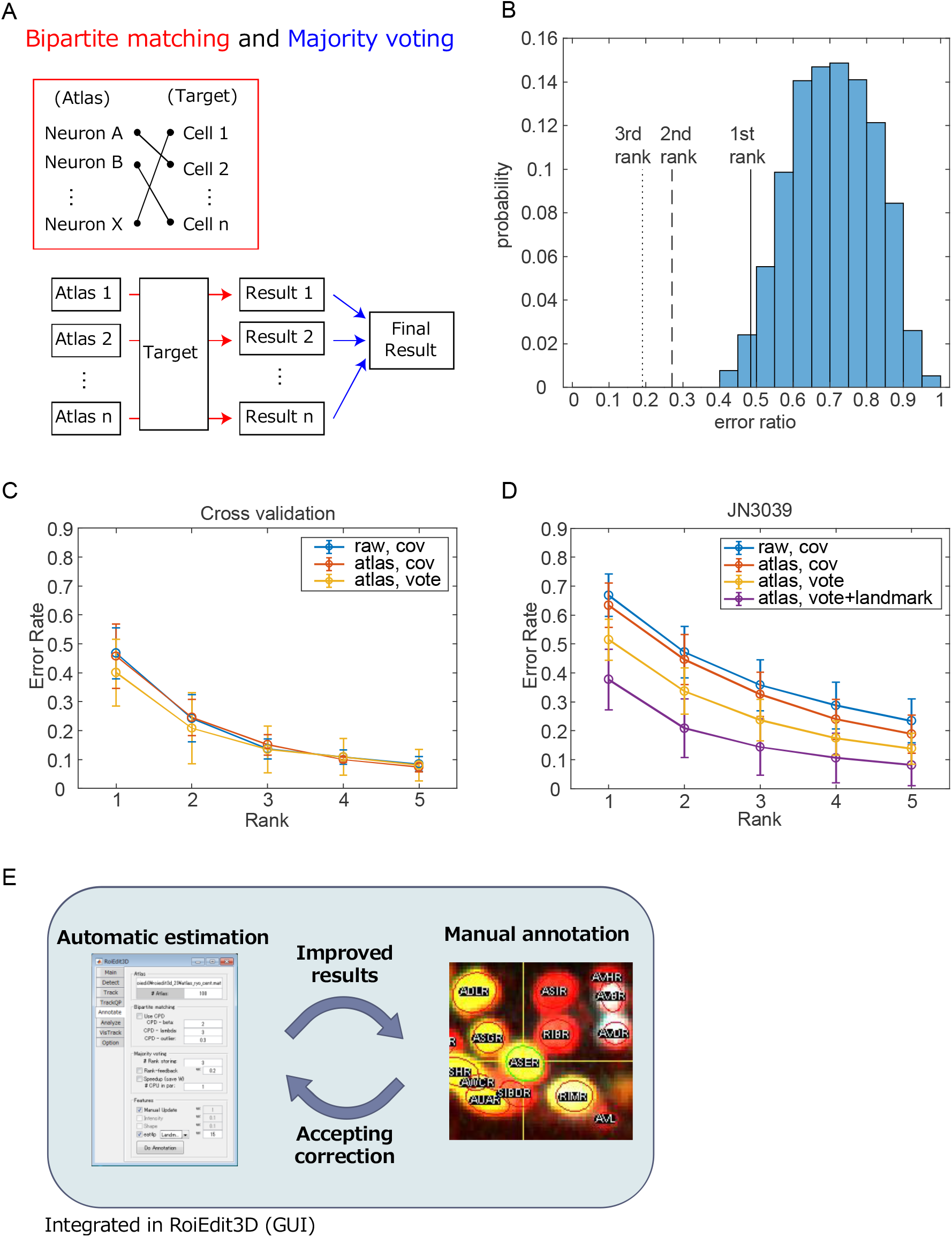
An automatic annotation method and evaluation. (A) The outline of the automatic annotation method. The schemes of bipartite graph matching and majority voting are shown. (B) Error rate of each bipartite matching and majority voting are shown in the blue histogram and the black lines, respectively. The rank N indicates that it is considered correct if the correct annotation appeared in the top N estimations. (C) Error rates of the automatic annotation method for the animals in the promoter dataset. The error rates were evaluated by cross-validation. (D) Error rates of the automatic annotation method for the strain JN3039 that expresses the fluorescent landmarks. (E) The automatic annotation method was integrated in the graphical user interface roiedit3d that enables feedback between automatic and manual annotation.

## Acknowledgements

A part of the computing resources was provided by the Shirokane3 supercomputing system of the Human Genome Center (the University of Tokyo). We thank the members of our laboratories for helpful discussions and technical assistance with the experiments. Information of neurons and expression patterns of promoters was provided by WormBase (www.wormbase.org) and by WormAtlas (www.wormatlas.org).

## Author Contributions

Performed the experiments: MK HS MSJ SO YM TT YT. Analyzed the data: YT SW MK HS MSJ SO YM TT YIw TI RY YIi. Wrote the paper: YT SW RY YIi. Conceived the project: YT YIi. Development and evaluation of the automatic annotation method: SW YT RY YIi. Designed the experiments: YT MK MSJ HS TI YIi.

## Figures and Figure legends

**Figure 1 - Figure Supplement 1:**
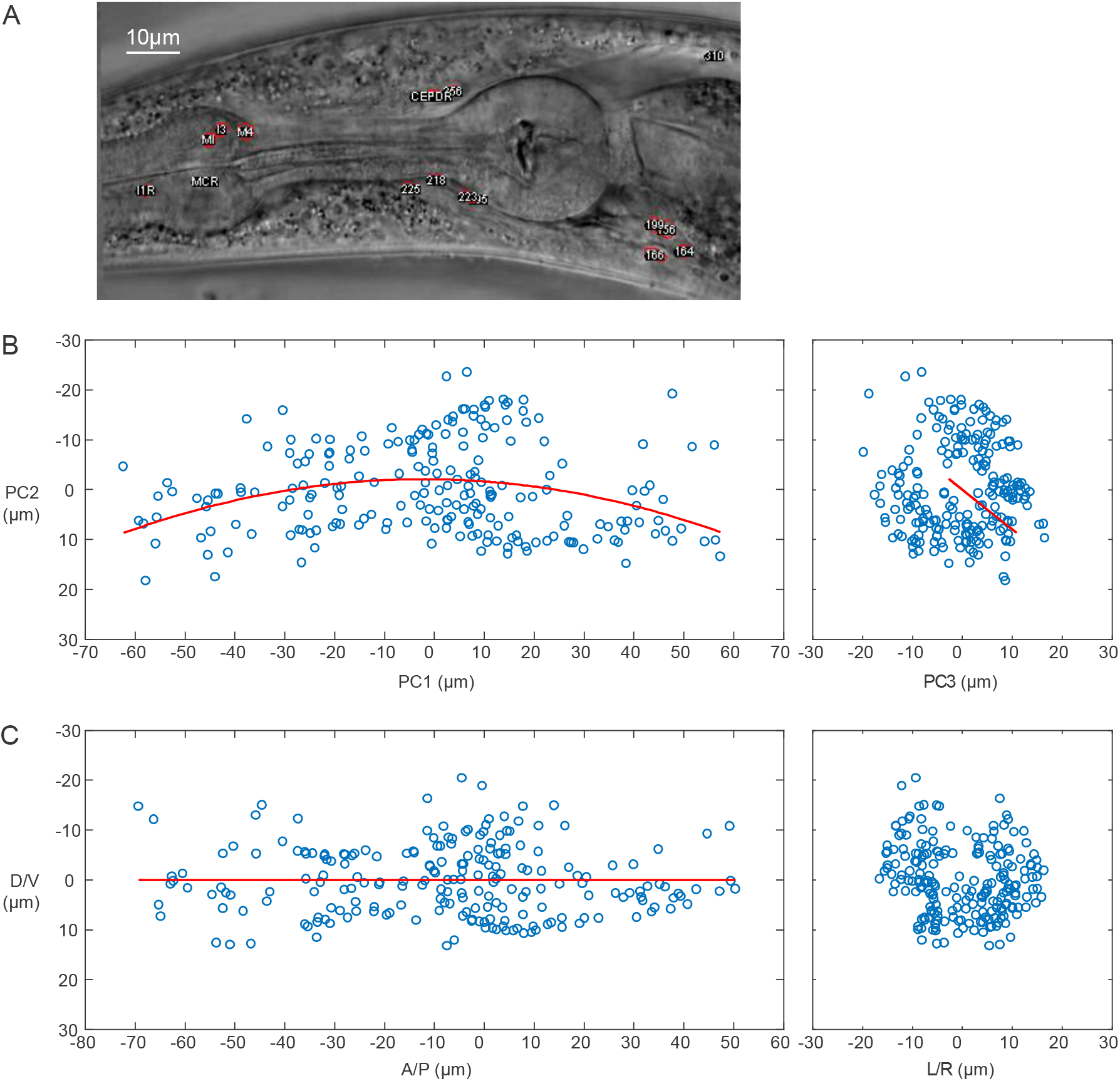
Correction of posture of the worms. (A) An example bright-field image of the head region of an adult animal with curved posture. (B) The positions of the cells in the animal (shown as blue circles) are projected onto the plane with PC1-PC2 axes and the plane with PC2-PC3 axes. The fitted quadratic curve is shown as the red line. (C) The corrected position of the cells.

**Figure 2 – Figure Supplement 1:**
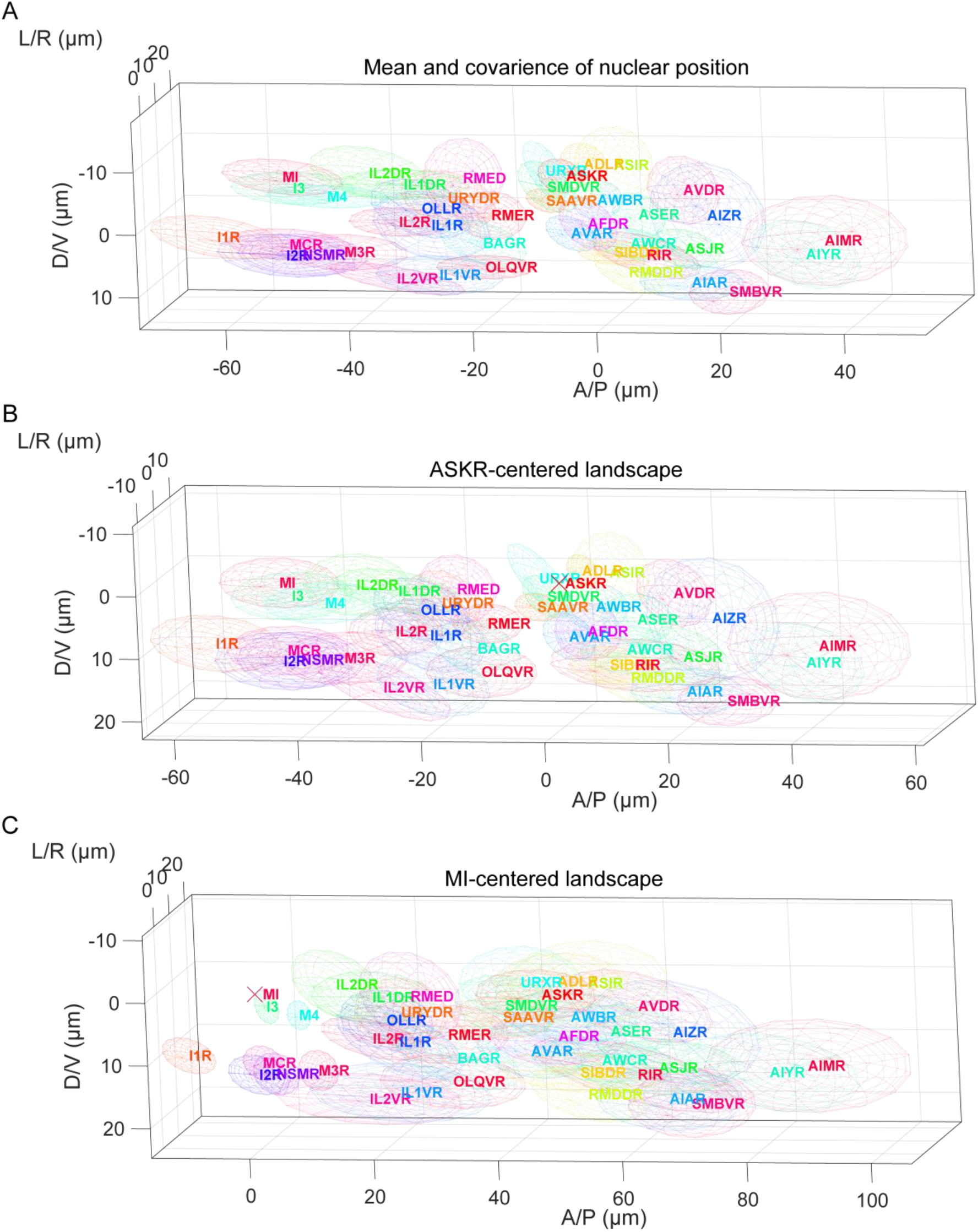
Specific-cell-centered landscape. (A) Original landscape as a reference. This panel is basically same as Figure 2A, but several cells are removed for visibility. (B) ASKR-centered landscape. The position of ASKR cell is indicated as a cross. (C) MI-centered landscape. The position of MI cell is indicated as a cross. The same cell have same color in (A)-(C). The cells in the right side are shown. Several cells are removed for visibility.

**Figure 2 – Figure Supplement 2:**
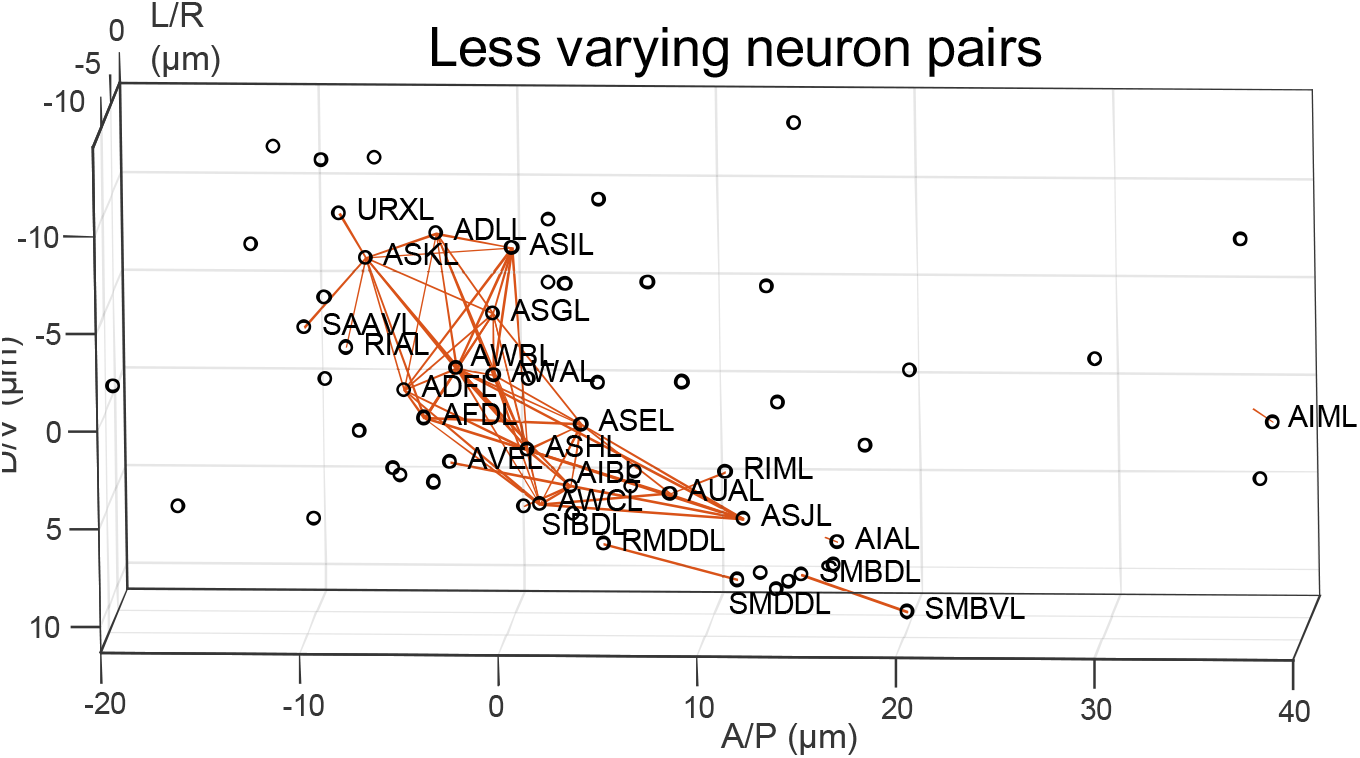
Less varying neuron pairs. Less varying neuron pairs were obtained by random permutation of animals (see Methods) and the less varying pairs in the left half are shown by red lines. The pairs including pharyngeal cells were omitted for visualization.

**Figure 2 - Figure Supplement 3:**
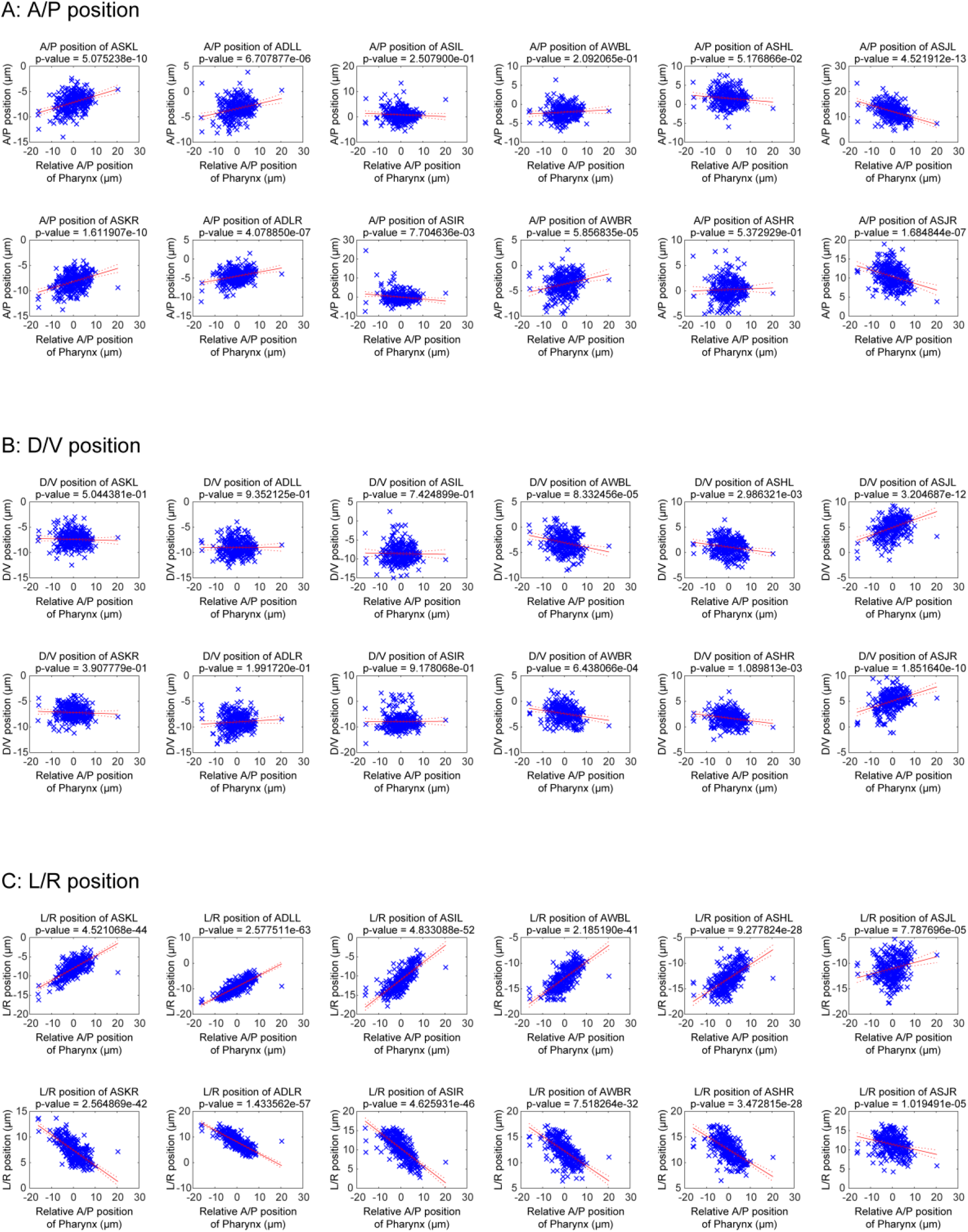
Position of posterior pharyngeal bulb affects cell positions. (A-C) A/P (A), D/V (B) and L/R (C) positions of dye positive cells with relative A/P positions of pharynx. The relative A/P positions of pharynx were calculated from mean difference of positions of pharyngeal cells from reference. Blue crosses indicate the cell positions in respective animals. The red lines and the red dotted lines indicate regression lines and 95% confidence bounds, respectively.

**Figure 4 - Figure Supplement 1:**
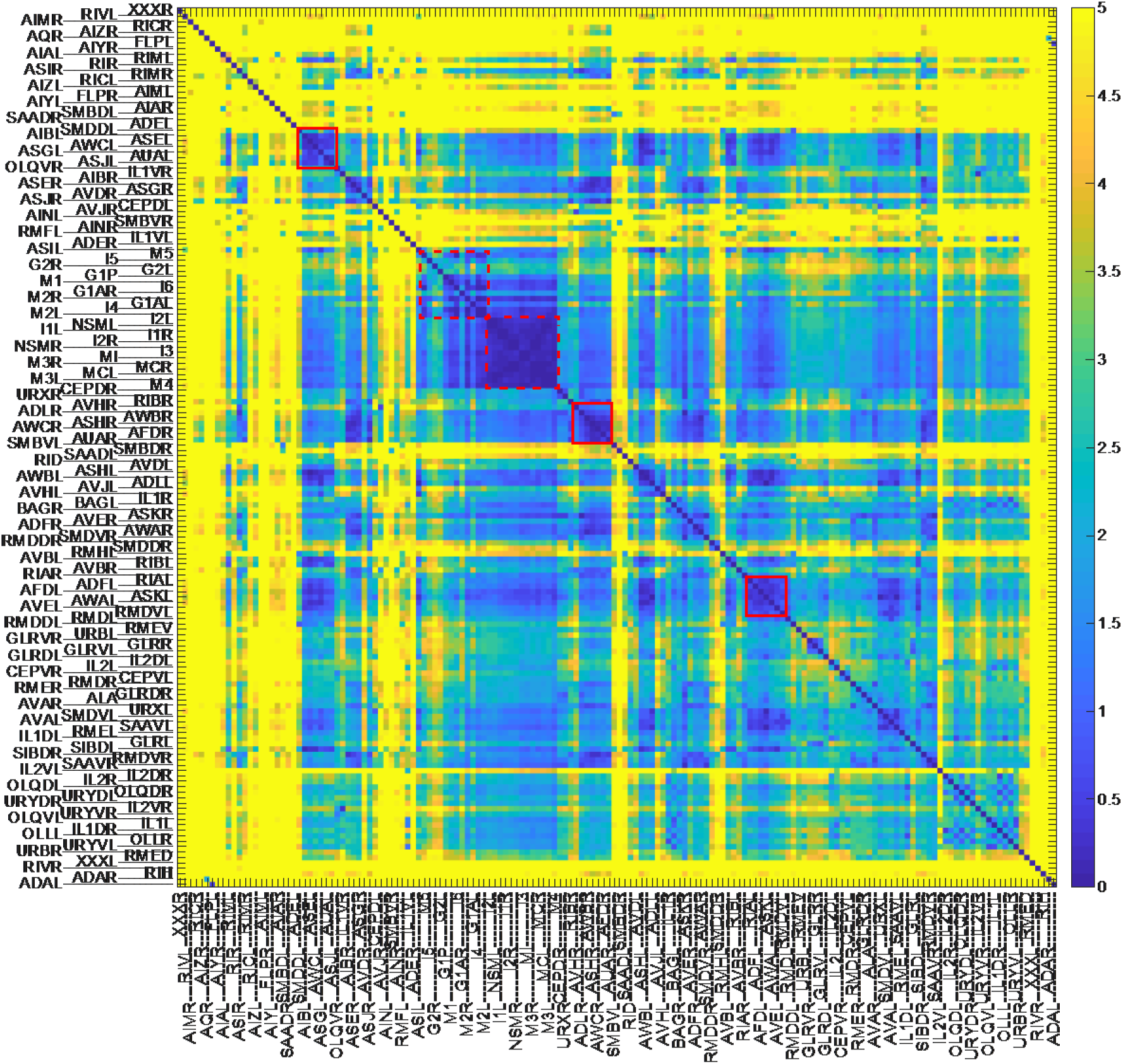
Variation of relative position of cell pairs. Orders of cells and colors are same as in Figure 2C.

**Figure 5 - Figure Supplement 1.**
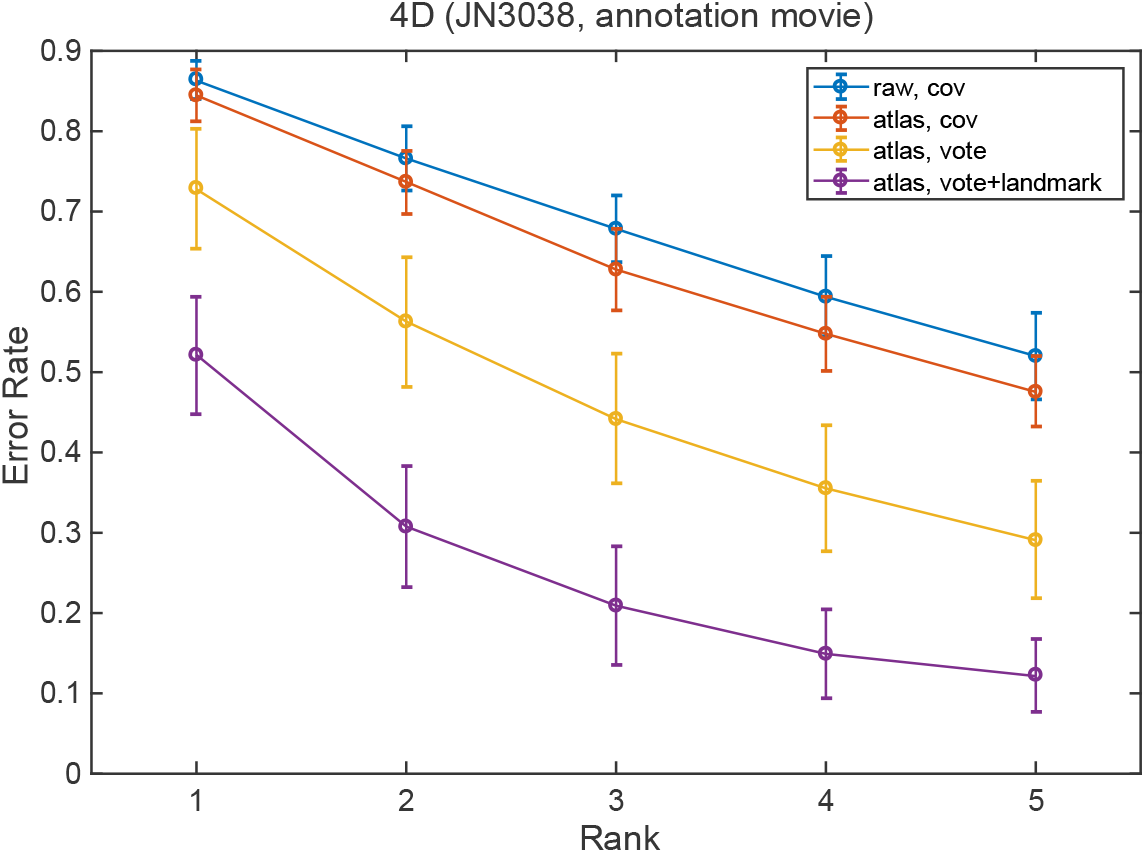
Error rates of the automatic annotation method for the animals in a microfluidic chip for whole-brain activity imaging (JN3038 strain, n=12).

## Supplementary Materials

- Supplementary Note 1: Optimization of parameters for atlas generation
- Supplementary Table 1: Evaluation result of promoter combinations (Excel file)
- Supplementary Dataset 1: Annotation dataset (contains positions and expression patterns) and corresponding static 3D images
- Supplementary Dataset 2: Positions of nuclei and expression patterns of landmark fluorescence in the whole-brain imaging strains as the test data for automatic annotation and corresponding static 3D images
- Supplementary Dataset 3: All codes for the GUI RoiEdit3D and analysis pipeline to make figures

All tables and datasets will be available from Figshare (10.6084/m9.figshare.8341088) upon publication of this paper. Current unpublished link: https://figshare.com/s/1e39bebd7568b41a39f5

