## Supplementary Note 1 for "An annotation dataset facilitates automatic annotation of whole-brain activity imaging of *C. elegans*"

Supplementary Note 1: Optimization for parameters in atlas generation

The generated reference samples serve as a set of virtual atlases that imitate observed topological variations of cellular positions across different worm samples. To obtain more realistic atlases, we optimized $\Sigma = \mathrm{diag}(\sigma_1, \sigma_2, \sigma_3)$ in Eq 1, which is the parameter to control the smoothness of displacements in the sequential alignments. First, we designed a measure $\boldsymbol{J_{t \to s}(i)}$ to characterize the coherency of the displacement of cell $i$ with respect to its neighboring cells in two images, $I_t$ and $I_s$:

$$\boldsymbol{J_{t \to s}(i) = \min_{r} \langle d_i - r, \bar{d}_{\mathcal{N}_i} - r \rangle \equiv \left\| d_i - \bar{d}_{M_i} \right\|^2}, \text{ where } \langle \cdot \rangle = \textbf{dot product}. \quad (2)$$

The displacement of cell $i$ was $\boldsymbol{d}_i = \boldsymbol{x}_i^s - \boldsymbol{x}_i^t$. The mean displacement of its neighbor set $M_i$ was calculated as $\bar{\boldsymbol{d}}_{M_i} = |M_i|^{-1} \sum_{j \in M_i} \boldsymbol{d}_j$ where $M_i$ denotes a set of cells neighboring within the squared distance less than 70 px with respect to cell $i$ in the image $I_t$. For each cell, this measure was calculated for all pairs of the 311 human annotation data if the number of neighboring cells was larger than four. In addition, we calculated $\boldsymbol{J_{t \to s}(i)}$ for randomly chosen 1,000 pairs of the computationally manipulated atlases. Finally, we used Bayesian optimization technique to minimize the Kullback-Liebler divergence between the normal distributions fitted to given $\boldsymbol{J_{t \to s}(i)}$ for the human annotation data and the computationally manipulated atlases, respectively.
